# Social licence through citizen science: A tool for marine conservation

**DOI:** 10.1101/266692

**Authors:** Rachel Kelly, Aysha Fleming, Gretta Pecl, Anett Richter, Aletta Bonn

**Affiliations:** Helmholtz-Center for Environmental Research – UFZ, Permoserstraße 15, 04318 Leipzig, Germany; German Centre for Integrative Biodiversity Research (iDiv), Deutscher Platz 5e, D-04103 Leipzig, Germany; Centre for Marine Socioecology, University of Tasmania, Hobart, Tasmania 7005, Australia; Institute for Marine & Antarctic Studies, Hobart, Tasmania 7001, Australia; Land and Water, CSIRO, Hobart, Tasmania 7001, Australia; Institute of Biodiversity, Friedrich Schiller University Jena, Dornburger Str. 159, 07743 Jena, Germany

## 1. Introduction

Meaningful public engagement with science and research is essential to improving knowledge about the environment and supporting sustainable use of ecosystems and natural resources. Transparent and culturally-appropriate natural resource management is imperative (Christie et al. 2017) and the public’s role in decision-making and data collection is increasingly recognised (Hecker et al. 2018), supported by a rise in the number of citizen science projects in recent years (Chandler et al. 2017;Pocock et al. 2017), including marine-based projects (Cigliano and Ballard 2017). In parallel, social licence has become an important theme for development, particularly towards exploring stakeholder engagement and communication (Lacey et al. 2017). Citizen science is often a partnership between the public and professional scientists to address questions and issues of common concern (Shirk and Bonney 2015) and has long been established as a means to engage society in science. Comparably to social licence, citizen science provides a means for stakeholders to have a voice in resource monitoring and in evaluating and promoting decision-making that might otherwise exclude them (Cigliano et al. 2015). This paper examines the potential of citizen science to enhance participant understanding of environmental sustainability and to promote social licence for nature conservation, examining marine citizen science as a case study.

The marine policy landscape is still young and emergent within Europe, and the European Union (EU) promotes sustainable growth of maritime and coastal activities as well as sustainable use of coastal and marine resources. Whilst initiatives have been adopted to enhance the protection of the European marine environment (i.e. Marine Strategy Framework Directive in 2008; Marine Spatial Planning Directive in 2014) some major challenges to the effective implementation of European marine environmental management and legislation remain. A large component of these include substantive criticism on ‘inadequate stakeholder engagement’ in EU policy making, and challenges hampering new and planned developments include a lack of social acceptance or social licence (see e.g. (Soma and Haggett 2015). Marine citizen science provides an interface between marine science and ocean literacy and offers a platform for the public to connect with science and policy and promote collaboration and partnerships between marine science and society. Consequently, in 2017 the European Marine Board (EMB) outlined a need to formally incorporate marine citizen science into EU research and policy (Garcia-Soto et al. 2017).

Development of citizen science in Europe is increasingly promoted and also supported by the establishment of the European Citizen Science Association (ECSA) in 2013 (Storksdieck et al. 2016) and the development of the ten principles of citizen science (Robinson et al. 2018). ECSA aims to promote sustainable development through citizen science, on the premise that environmental sustainability and stewardship are intrinsically linked to research, innovation and empowerment (Storksdieck et al. 2016). The expansion of European citizen science has also seen increases in the number and diversity of marine projects (Garcia-Soto et al. 2017). Community networks can play a strong role in enhancing policy support (Dean et al. 2016) and citizen science presents an ideal tool to explore this. Citizen science is known to engage and inform the public about science and the natural environment and enhance empowerment to act (Martin et al. 2016a;McKinley et al. 2017;Nursey-Bray et al. 2017). Such public engagement can provide an avenue to develop social acceptance, allowing communities and society to partake in policy development and decisions that will affect them (Soma and Haggett 2015).

Citizen science can be a means to foster collective action, by providing opportunities for individuals to participate in coordinated research efforts (Shirk et al. 2012). Participants can provide direct input into decision-making by using knowledge learned through citizen science to comment on policy action. They can also indirectly affect policy, by disseminating information amongst their communities, educating and motivating others to become involved in natural resource conservation and policy discussion (McKinley et al. 2017). Citizen science helps to empower the public by promoting learning-by-doing (Bela et al. 2016), providing opportunities for true participation in science and evidence provision for decision-making, and encouraging them to adopt more active roles in society (EuropeanCommission et al. 2013). It can enhance flows and exchange of information between communities, scientists, marine managers, and policy decision-makers to help produce solutions that promote better environmental and social outcomes and contribute to conflict mitigation in natural resource management (McKinley et al. 2017). Further, it is known that the greater the involvement of local people in monitoring activities, the quicker it is for this data to be implemented into decision-making (Danielsen et al. 2010).

Citizen science programmes are rapidly gaining acceptance as an integral part of society, science and policy (Pecl et al. 2015), but large gaps remain in our understanding of the utility of citizen science in a marine policy and management context. This paper is among the first attempts to link social licence theory with citizen science, aiming to produce practical outcomes that can be applied to sustainable ocean management. Firstly, we explore the concept of social licence and how it might relate to citizen science (Table 3). Then, we examine marine citizen science in Europe, through survey and interview methods, to explore its potential role in creating social licence. We identify how these two concepts influence knowledge and opinions and connect diverse users of the marine environment to promote engagement, ocean literacy and trust in European marine conservation.

### 1.1 Social licence and the European marine context

Social *licence* suggests that society (i.e. communities and stakeholders) can award and withhold some kind of permission for an activity. Whilst of no direct legal value, the concept owes its considerable power to the legal ramifications it can indirectly incur on resource users. Social licence is theorised as ongoing acceptance or approval from stakeholder communities, a process that requires establishing meaningful partnerships between operations, communities and government based on mutual trust (Parsons and Moffat 2014). No concise definition of social licence exists, nor does this paper seek to achieve one. This study aims to determine novel means through which social licence can be developed to improve sustainability in the marine environment. Gaining social licence implies achieving and maintaining public trust that resource users and managers are utilising marine spaces and resources ethically, in accordance with societal expectations.

It is increasingly evident that social licence is important for using, developing and protecting marine spaces but it remains unclear how social licence might best be achieved through public engagement in practice. Whilst some research has been initiated on social licence for the ‘Blue Economy’ (Soma and Haggett 2015), to date and to our knowledge, no research has been conducted into social licence for biodiversity conservation or specifically, marine conservation in Europe. Understanding social acceptability is crucial for marine management (Gall and Rodwell 2016). Failure to incorporate social and cultural considerations into natural resource conservation initiatives can result in significant conflict and negative environmental outcomes.

If the emphasis of the concept of social licence is on societal benefits and transparent decision-making processes (Smits et al. 2017), then public engagement can shape social licence and increase accountability of decision-making (Soma and Haggett 2015). However, it cannot be assumed that simply engaging the public will result in trust, support and acceptance; more meaningful and in-depth relationships between user groups and the ocean environment are required (Mercer-Mapstone et al. 2017). Building advocacy through citizen science may foster social licence for natural resource protection and management because partnerships achieved through citizen science can create pathways for support between user groups and networks for collaborative decision-making. Stakeholder analysis is relevant to conservation research because management impacts likely involve different uses and user groups and intersect both biological and social systems (Brown et al. 2016). We do not explore this here, but highlight stakeholder analysis as a pertinent area of research. Multiple definitions of ‘stakeholders’ exist in the literature, in this study however we adopt Grimble and Wellard’s (1997) definition:

> *“…any group of people, organized or unorganized, who share a common interest or stake in a particular issue or system … who can be at any level or position in society, from global, national and regional concerns down to the level of household or intra-household, and be groups of any size or aggregation” ((Grimble and Wellard 1997) p. 175)*

Whilst stakeholder definitions vary depending on context and situation, there is widespread agreement on the need and importance of incorporating these groups into marine conservation management through meaningful participation and engagement (i.e. (Voyer et al. 2012;Brown et al. 2016). Both local and non-local engagement are important in the pursuit of social licence for marine conservation (Munro et al. 2017). Citizen science is ideally placed for this engagement and can act as catalyst for individual behaviour change that is linked to environmental stewardship of marine systems (Cigliano et al. 2015).

Participation in citizen science can instill volunteers with a sense of ownership, both of the data they collect (Reed 2008) and the areas that they monitor (Newman et al. 2017). Citizen science may also be valuable in promoting scientific (ocean) literacy amongst the public (Bonney et al. 2009). With regards to the marine environment, the EMB has highlighted ocean literacy as a strategic action area for improving public awareness on the value of marine environments and research and to promote interest and participation amongst civil society in marine policy (EuropeanMarineBoard 2017). In this paper, we identify how marine citizen science may influence knowledge and opinions, connect diverse users of the marine environment and improve ocean literacy to promote social licence for marine conservation in Europe and potentially elsewhere.

## 2. Methods

We used an adaptive theory approach in this study, akin to Vann-Sander et al. (2016), because comparing and contrasting social licence and citizen science was anticipated to generate new theory and it was necessary to ensure that all relevant information on the topic was captured effectively as it emerged through the research process. For a thorough examination, we chose to combine in depth qualitative interviews with an online semi-quantitative survey with marine citizen science managers. We adopted this mixed-methods approach so that we might engage as deeply and actively with participants as possible, to understand their perceptions of marine citizen science, and its connection to social licence occurring in practice, whether in their opinion it was successful and how it might be potentially useful for generating social licence in the marine space.

### 2.1 Surveys

The initial research phase consisted of an online survey of marine citizen science project coordinators to obtain information on the extent of projects in Europe and their aims (i.e. education, data collection), as well as coordinators’ perceptions of European marine management and conservation. The sample of projects was obtained from the EMB (Garcia-Soto et al. 2017) and was further supplemented by sharing the survey online amongst colleagues in wider networks to disseminate it to other potential respondents. Of the initial n=60 project coordinators approached, 35 (58.3%) coordinators responded to the online survey (conducted using Lime Survey GmbH).

### 2.2 Interviews

Following the survey, potential interviewees were identified from the online survey respondents. All respondents were invited to partake in the interview stage and 15 agreed to do so. These semi-structured interviews were conducted by the lead author in July, August and September 2017. The interviews lasted between 30-80 minutes, were conducted over the phone, and were audio recorded before being transcribed by the lead author. The interview questions focused on the organisation of citizen science, project objectives, their development and potential connection to social licence. A full list of survey and interview questions is provided in Appendix A.

All interview transcripts were subject to thematic analytical evaluation using NVIVO (for Mac Version 11.4.2-2081). Initial codes were generated and themes were developed iteratively using a grounded theory approach (Haywood 2015). These were reviewed, compared and redefined where necessary and relationships between codes were identified. Hierarchical coding was used to organize themes and to identify the resulting six key themes of the study. These themes are presented below and represent the aggregated interview responses, as opposed to the questions that guided them.

## 3. Results & Discussion

The online survey responses represent 34 projects of varied sizes and purposes from more than 8 different European countries, the majority of which are located in the U.K. (19 projects or 55.8 %). A full list of projects, locations, focus and participation can be seen in Appendix B. The coordinators’ responses represent a diversity of projects, but not a diversity of countries and are largely pooled from English-speaking countries. We highlight this possible language barrier as one that might be addressed in future studies. The surveyed projects vary considerably in size, scope and intent. Project design will influence its potential to collect and share scientific information and engage with the public (Shirk et al. 2012). In agreement with other studies on citizen science, the projects described here generally do not formally document and report on any participant learning objectives or achievements (Bela et al. 2016).

**Table 1.**
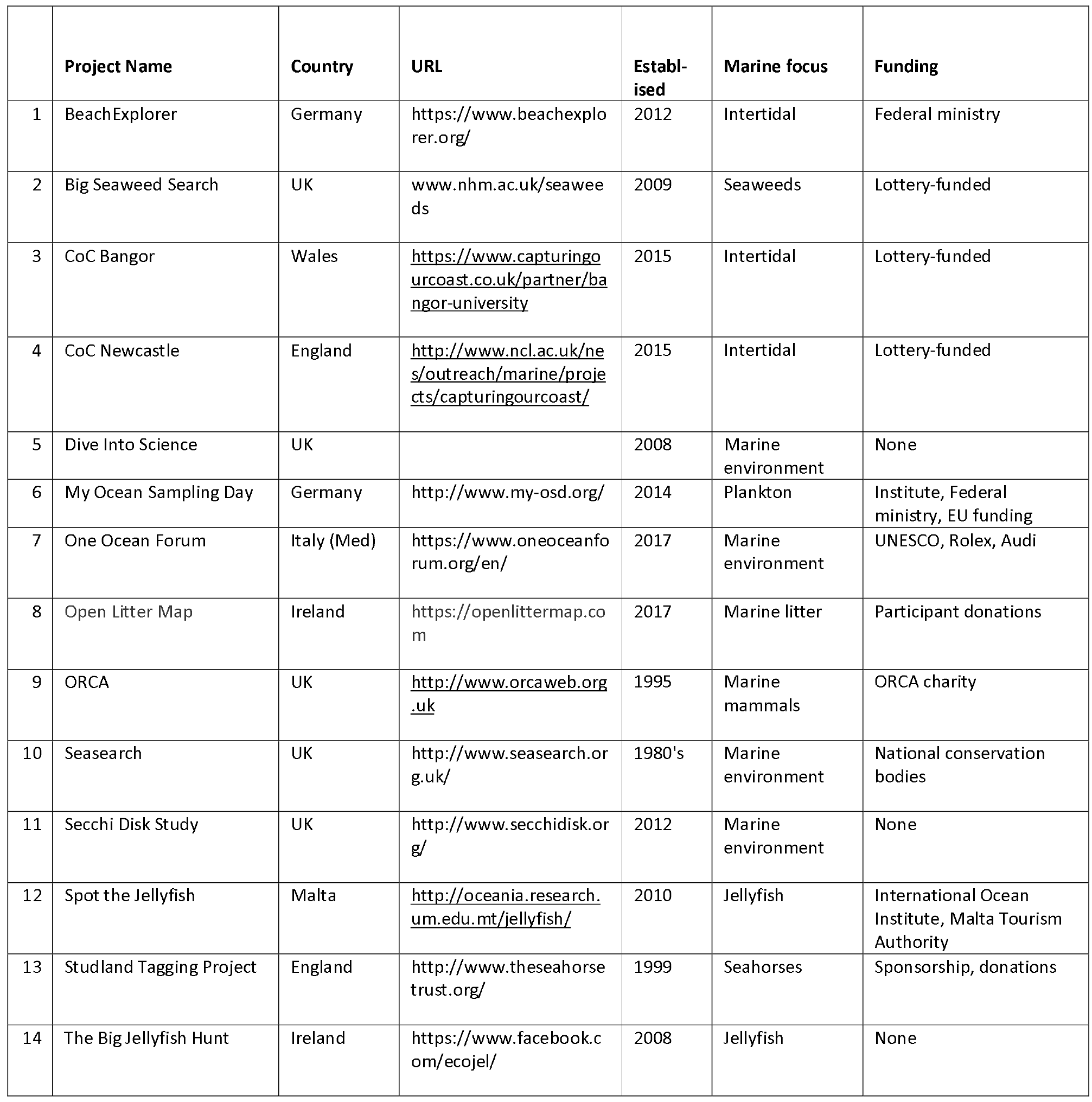
Overview of the 15 marine citizen science projects engaged in interviews. These projects represented 6 countries, had varying levels of establishment (i.e. long-established, to very new projects), different objectives in regards to marine research and conservation, and were supported by different schemes.

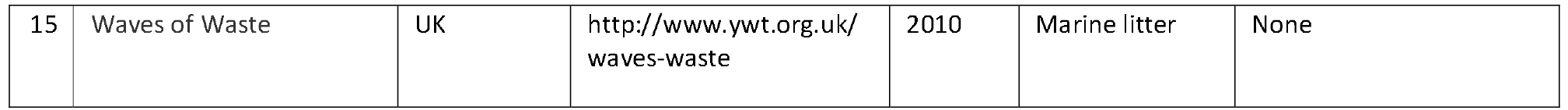

The interview coding produced five key themes: *developing understanding, communicating, logistics, future of citizen science, people, and connecting. Developing understanding* was the most commonly identified theme, with 147 references across all 15 sources. Connecting was the least mentioned, with 76 references across all 15 sources (Table 1). Below, we elucidate these five themes and refer to existing literature (Table 3) to identify their interactions with, and roles in, citizen science and social licence.

**Table 2.**
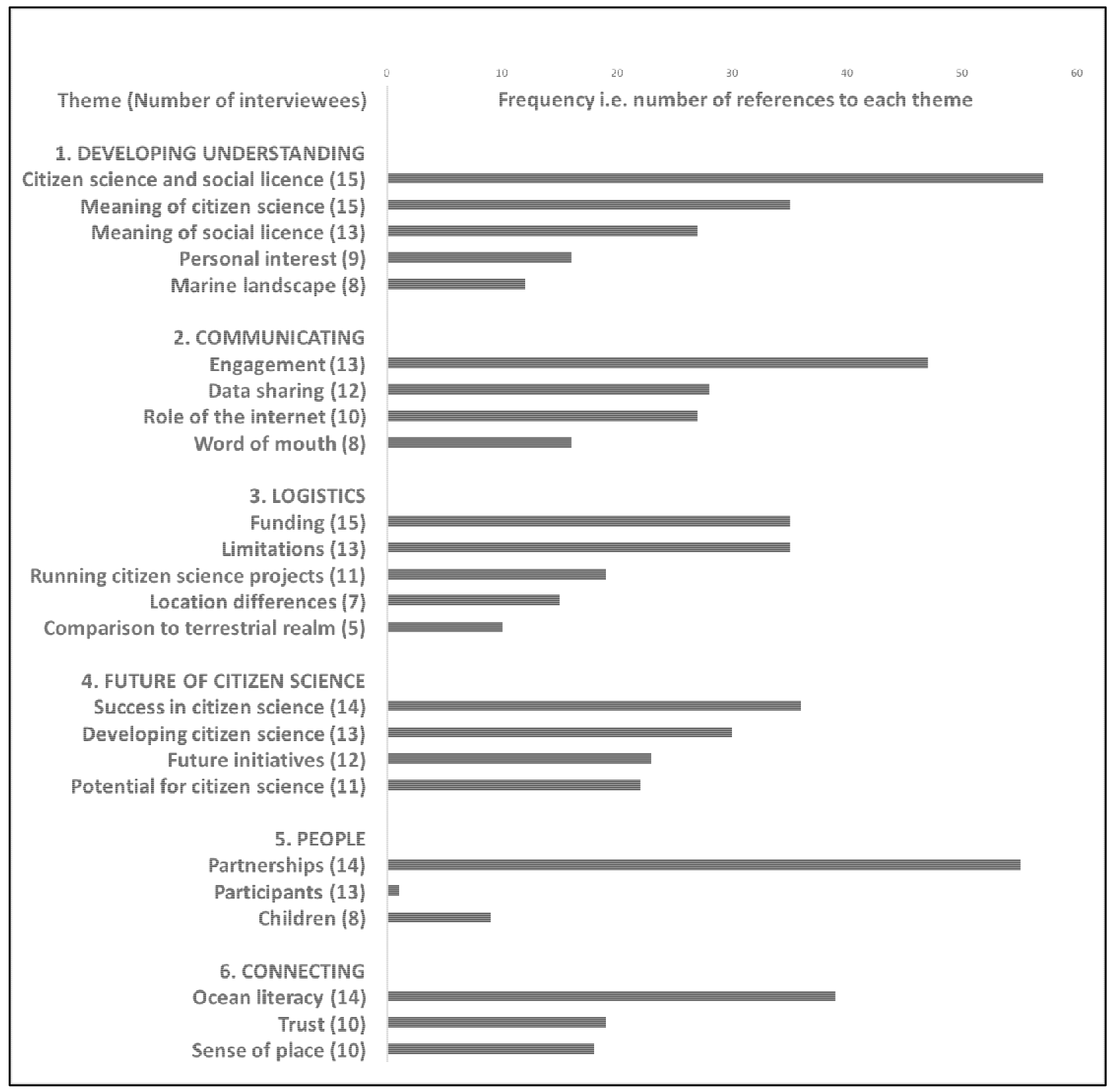
Interview codes Note: the frequency (i.e. number of references) indicates how frequently each theme was mentioned in interviews, rather than the importance interviewees placed on these issues.

### 3.2.1 Developing understanding

This theme largely focused on developing understanding about the concepts of citizen science and social licence. There was strong support in favour of using citizen science to generate social licence for marine conservation. It was widely accepted that creating social licence would require specific project design and objectives and also frequently highlighted that *‘the first step in that is people have to care and be engaged with that kind of environment and citizen science definitely builds that sense of ownership…’(C2)*. It was generally felt that social licence actions were already happening to some degree in many places, e.g. a petition to legally protect seahorses, community resistance to coastal development plans. However, the actual term ‘social licence’ was new to all but one of the interviewees. Many participants, particularly from the UK and Ireland, synonymised it to ‘buy-in’; ‘public acceptance’ and ‘public pressure’ were also terms used amongst interviewees. Coordinators’ accounts and understanding of the role of citizen science and social licence largely tied into themes discussed in other studies, i.e. citizen science can enhance scientific literacy, improve ecological knowledge, promote connections with nature and locality, strengthen social ties, and influence participants’ sense of stewardship and the responsibility they feel towards the environment (Haywood 2015):

> *‘It comes back to the simple thing of bridging the gap and making them feel valued and having an important role marine conservation, which is what citizen science does, it gives them that buy-in’(A4)*.

Interviewees’ understanding of the term ‘citizen science’ varied depending on the context or scope of their project. Terminology is particularly dynamic in citizen science because the field continues to develop, expand and diversify (Eitzel et al. 2017). Most coordinators did not wish to be restricted by definition and were keen to extend their projects to broader groups and to partner with other schemes that did not necessarily conduct citizen science. One coordinator did, however, take umbrage with the term ‘citizen science’ and preferred to use ‘conservation volunteers’, a term that he found was more accepted by, and accessible to, his project’s participants. Certainly, the meaning of citizen science can represent different things to different people and create confusion about its nature and utility (McKinley et al. 2017). We highlight that one of the challenges of using citizen science as a tool to create social licence, is that the objectives of citizen science need to be transparent to participants (see ‘Cooperation’ in Table 3). Defining such objectives with participants can be considered a project objective in itself.

### 3.2.2 Communicating

The theme of communicating largely focused on engagement, the different means by which marine citizen science projects interacted with their participants and how participants shared this information more widely. Engagement and sharing knowledge about the marine environment was seen as a *‘very strong purpose’(C2)* of marine citizen science. Modes and frequency of engagement varied widely (i.e. newsletters, seminars, beach-meets, training sessions, online forums, email updates, beachside billboards, etc.) and were shared frequently (often daily) to very rarely (largely due to funding or time constraints). Consistent with other studies, coordinators highlighted the value of personal and face-to-face communication with participants in developing rapport and for engendering meaningful relationships beyond transactional interactions (Martin et al. 2016a).

There was a large consensus that *‘communication is key’(A4)*. Many coordinators underscored the role of the internet in their ability to share information and communicate more efficiently with a wide public network and more easily for both participants and organisers. Social media (i.e. Facebook, Twitter) improved projects’ ability to recruit participants and to remain in contact with them. For example, The Big Jellyfish Hunt, is a project that communicates to its participants only through Facebook; Open Litter Map, one of the youngest projects in this study (established 2017), is also only internet-based. The importance of the internet for these projects is not surprising. Mobile technologies have moved society beyond its traditional infrastructure barriers, facilitating much broader participation in citizen science programmes that make use of developing technologies (Pimm et al. 2015). However, different marine user groups require different engagement strategies and projects must consider their own goals and capacities when designing and implementing participant engagement (Hind-Ozan et al. 2017). For instance, social licence is founded on meaningful dialogue and communication (Yates and Horvath 2013), but exploration is required as to whether citizen science can best achieve this through face-to-face or digital media interactions.

Data-sharing was an objective for several of the projects, particularly those who developed partnerships with government or academic institutions. Many projects provided data used in marine protected area (MPA) designations and now contribute to monitoring efforts within these MPAs. Others published the data in scientific papers in peer-reviewed journals. The ORCA Trust is the lead partner of the European Marine Cetacean Monitoring Coalition, a consortium of eight cetacean monitoring organisations across Europe that are *‘collecting data to help inform policy and legislation, to improve the conservation of our marine space’(A4)*.

Sharing of data was seen as a major influencing tool for marine citizen science. It was agreed that *‘people spreading the word’(B1)* on data they collected or new knowledge they learned through marine citizen science played a big role in disseminating information to the wider public (i.e. participants’ families, friends and community networks). These observations align with other experiences in the literature which show that volunteering in citizen science projects increases participants’ concern about conservation issues and that participants disseminate the knowledge they learn to their wider social networks (Johnson et al. 2014;Nursey-Bray et al. 2017). Citizen science can have broader societal impacts, especially in promoting conservation awareness because *‘personal conversation is probably the biggest spreader of education’(B4)*. In Table 3 below, we outline how citizen science also promotes reflection and discussion on how science interacts with society and societal values and how we can embed this more deeply into public thinking and decision-making (Storksdieck et al. 2016).

Similarly to social licence, citizen science provides a means through which communities and stakeholder groups can participate and have a voice in marine resource management and decision-maing, that might otherwise exclude them (Cigliano et al. 2015). Involving citizens in data collection that will inform marine management can legitimise data and generate trust in its validity and application. ‘Marine citizenship’ (i.e. an individual’s rights and responsibility as regards the marine environment) necessitates increased awareness about marine issues, adequate understanding about the personal role and behaviour involved in creating and solving these issues and a positive shift in values that can promote ocean-friendly, pro-environmental behavioural decisions (McKinley and Fletcher 2012).

### 3.2.3 Project logistics

Funding was identified as a limitation to development and engagement by most coordinators and is a pertinent issue that must be addressed if marine citizen science is to benefit marine conservation in Europe. Citizen science can be a cost-effective means to gather data for scientific research (Aceves-Bueno et al. 2015) and we highlight the multiple benefits of investing in citizen science development, to enhance both scientific, social and political outcomes (Hecker et al. 2018) for European marine conservation. Improved funding can also enhance the likelihood of producing accurate and pertinent data for marine conservation. The funding sources that supported projects in this study varied greatly and included government grants, corporate sponsorships, scientific institutes, lottery funding, donations, membership fees, amongst others. Several projects had no direct source of funding whatsoever. Unsurprisingly, these projects lacking funding struggled to expand their engagement, recruitment and research activities.

Other limitations included meeting participants’ expectations and incorporating diverse values into developments and partnership-building: *‘There’s a lot of politics in conservation, as I’m sure you’re finding out, that stops a lot of partnerships’(B4);* and successfully retaining participants that were recruited to projects: *‘That is always a challenge, how do we get more people interested?’(C5)*. These limitations further emphasise the need to increase availability of specific resources for marine citizen science projects, to enhance potential partnerships and promote public engagement. Investing in marine citizen science can enhance project capacity to engage more widely with the public, address societal concerns in its research, and promote resulting data to both communities and decision-makers.

### 3.2.4 Future of citizen science

Developing marine citizen science projects was a strong subject in this theme. Whilst several of the projects were stagnant as a result of funding or other constraints, and others were only becoming established, all projects were hoping to develop and even expand their scientific activities and engagement. Coordinators emphasised that marine citizen science is *‘not a one-size-fits-all approach’(A5)* and that two-way communication between participants and coordinators is vital for developing projects that can be maintained successfully in the long-term. In advancing marine citizen science for the value of science and policy, planners must be careful to match their programmes’ methods of engagement, public involvement and participation appropriately with their type the project and its focal aims (McKinley et al. 2017). Table 3 (below) serves to juxtapose social licence and citizen science, so that planners may identify how and where to best enhance social licence, i.e. for marine conservation through citizen science activities.

The coordinators discussed their project successes in educating and improving people’s understanding of marine species and ocean environments, particularly the success of marine citizen science in promoting ocean literacy: *‘They always learn something new, they always get excited’(A2)*. The majority of coordinators spoke of their very positive experiences of participant education and improved awareness and how this had changed and enhanced participants’ perceptions of the marine environment. However, several did articulate concerns on whether citizen science projects outcomes have the potential to reach all members of the public, and the difficulties in retaining participants for longer time periods. These challenges are also felt in social licence issues, where some members of the public are more engaged than others and where the ‘loudest voice’ might not in fact be the most representative (Cullen-Knox et al. 2017). An objective for future citizen science and social licence research may be in determining how to ignite and sustain interest and care for marine science and conservation issues (Ballard et al. 2018).

**Table 3.** Synthesis of key elements common to social licence and citizen science. We identify common elements described in the literature and identify relatedness between the concept of social licence and practice of citizen science.

| **Social Licence (SLO)** |  | **Citizen Science (CS)** |
| --- | --- | --- |
| Communicating and constructing two-way, meaningful dialogue with and between stakeholders is essential to generate SLO (Moffat and Zhang 2014; Zhang et al. 2018). | **Engagement (Relationships)** | Maintaining volunteer participation in CS is required to build and develop the capacity of any project (Bonney et al. 2009; Martin et al. 2016b; Nursey-Bray et al. 2017). |
| Addressing SLO issues allows local communities to raise concerns and opinions they might not otherwise have the opportunity to share (Edwards and Lacey 2014; Jijelava and Vanclay 2017). | **Representation (Stakeholder voices)** | Developing CS collaboratively between researchers and the public can address issues and questions of community interest and enhance societal relevance of science (Thiel et al. 2014; Bonney et al. 2016). |
| Earning SLO requires bringing stakeholders together to discuss, debate and define issues and improve community relations (Moffat and Zhang 2014; Lacey et al. 2017). | **Connecting stakeholders** | CS brings diverse users together to share information and experiences, building relationships that might otherwise not exist (Aceves-Bueno et al. 2015; Bonney et al. 2016). |
| Communicating and debating groups' interests and concerns encourages dialogue and cooperation to achieve agreement and earn SLO (Gallois et al. 2016; Zhang et al. 2018). | **Cooperation (between groups)** | Working together in CS brings scientists and non-scientists to develop and achieve joint research and educational objectives (Bonney et al. 2009; Nursey-Bray et al. 2017). |
| Sharing perceptions, opinions and experiences can enlighten stakeholders, industry and government on the experiences of other groups (Gallois et al. 2016; Jijelava and Vanclay 2017). | **Education (Scientific literacy)** | Learning-by-doing in CS enhances understanding and scientific literacy. Participants also gain greater awareness about threats to their examined ecosystem through direct experience (Bonney et al. 2009; Crall et al. 2012). |
| Earning SLO demands that parties demonstrate that their use of the ecosystem and data is credible, legitimate and trustworthy (Moffat and Zhang 2014; Gall and Rodwell 2016; Jijelava and Vanclay 2017). | **Legitimacy (of science)** | Citizen science promotes reflection and discussion on how science interacts with society and its values. Jointly developing projects legitimises data collection, production and application (Aceves-Bueno et al. 2015; Göbel et al. 2016; Elliott et al. 2017). |
| SLO gives a voice to communities, that they might act as overseers of their local environments and resources (Boutilier 2014; Cullen-Knox et al. 2017). | **Stewardship** | Connecting the public to the natural environment through CS can increase awareness, attachment and willingness to protect it (Danielsen et al. 2010; Crall et al. 2012; Chen et al. 2015; Bonney et al. 2016; Newman et al. 2017). |
| Legitimising research and/or use of the natural environment increases the trustworthiness of the decision-making it informs (Boutilier 2014; Jijelava and Vanclay 2017). | **Trust (in decision-making)** | Promoting public engagement in collecting evidence that informs management increases understanding and trust in these management interventions (Aceves-Bueno et al. 2015; Hind-Ozan et al. 2017). |

### 3.2.5 People

Partnership-building with other groups and organisations was a means for projects to *‘strengthen both the research data, the quality of the data we were getting and the engagement and messaging we were doing’(C2)*. Citizen science can bring experts and non-experts together in partnerships that foster shared positive action in co-creating knowledge and information (Dickinson et al. 2012;Jordan et al. 2012). Whilst levels of involvement and influence varied, benefits that projects sought and gained through partnerships included the ability to recruit more participants, more scope to engage with the public, enhanced ability to share data they collected and larger pools of funding to expand their projects’ activities. In the UK in particular, most projects were affiliated with universities and government agencies, likely reflecting the long-standing tradition and development of citizen science in the British Isles. Coordinators believed that the larger their project network, the larger the impact their projects activities could have:

> *‘The larger the diversity with citizen science I think, that the higher are the chances it has an impact on social licence’(C4).*

Interestingly, participant types varied across and within projects, recruiting from *‘every single walk of life, from dustbin men to scientists to all those in between’(B4)*. This is consistent with a growing body of literature that recognises citizen science participants as diverse and representative of many kinds of people (Cigliano et al. 2015). Again, reflecting the value of marine citizen science for engaging with a large body of the European public, enhancing their ocean literacy and promoting social licence for marine conservation effort. However, other research has demonstrated that a large proportion of marine citizen science participants are more highly educated that the general public (Martin et al. 2016b). This is certainly a pertinent area for exploration that would guide development of recruitment and engagement, for both marine citizen science in Europe and elsewhere. It is important to consider *who* participates in these projects when developing marine citizen science to enhance ocean literacy and promote social licence. Where developed appropriately, the participatory structure of citizen science can certainly promote inclusion of diverse perspectives in decision-making processes (McKinley et al. 2017) as well as increase the legitimacy and social licence of decisions made for the marine management.

### 3.2.6 Connecting

The interviewees largely agreed that marine citizen science provides a valuable platform on which to educate the public about the ocean and also to connect them to marine environments they would not normally be aware of, or have exposure to. Participation in marine citizen science was considered a pivotal step for generating ocean literacy and reducing the *‘disconnect between people and nature’(C5)*. However, there was consensus that developing marine citizen science for this purpose would require adequate planning to address these objectives. The process of earning social licence is similar to citizen science because it brings members of the public together to discuss and address issues of common concern. The opinions of the coordinators agree with reflections in the literature on the need to properly understand the potential of citizen science as a communication and engagement tool (Groulx et al. 2017).

Citizen science is undoubtedly a valuable component of environmental stewardship (McKinley et al. 2017) because participants most frequently have strong positive attitudes towards the environment, demonstrate pro-environmental behavior and believe that their actions contribute to the value of natural resource conservation (Merenlender et al. 2016). Stewardship also plays a role in social licence because it gives communities a ‘voice’ to oversee usage and development of their local environments and can instil public resonsibility for natural resources (Table 3). Projects in this study demonstrated that *‘citizen science gives them [participants] a closer relationship with their local environment, or whatever environment they’re sampling from…ultimately gives people a greater understanding of the natural world and the environment in general’(B5)*. These feelings of connectedness and ownership are known to increase participants’ trust in the citizen science they are contributing to (Dickinson et al. 2012). These feelings of trust are a major determinant in whether participants will award social licence or not (Boutilier 2014).

Marine citizen science was seen to legitimise science and marine conservation by connecting people to their local and/or marine environments and generating a sense of place through ownership of that space:

> *‘It’s more likely that people protect what they know and what they value’(A3);*
>
> *‘It gives ownership to an area to stakeholders who normally feel disconnected’(B3);*
>
> *‘I think that is very, very powerful, when you get the locals themselves caring about the marine environment’(B1).*

This is in agreement with other studies who have shown that people frequently need to personally experience the ocean (and its problems) before they are likely to change their views and attitudes (Steel et al. 2005). Leveraging this ‘power of place’ is posited as a valuable means to improve conservation decision-making and increase participation in citizen science (Newman et al. 2017). We identify this sense of place component as one that requires future exploration and development.

Trust was also an important topic under this theme. Likewise, this relates to communication because participants who continue to be engaged effectively will continue to trust citizen science projects and outcomes (Hind-Ozan et al. 2017). Further, citizen science data can educate already pro-environmental participants and help them disseminate and argue the importance of marine conservation amongst their wider networks (Cigliano et al. 2015). This can legitimise research and the data collected and increase the trustworthiness and social licence of the marine management decisions it informs (Table 3). The project coordinators largely agreed that developing trust for marine conservation in Europe is a complex challenge that will need to be met with complex, complimenting approaches because often, *‘people trust what they want to hear’(C4)*. Ownership, developing ocean literacy and marine stewardship were seen as requirements for generating understanding and personal connection to the ocean and trust in decision-makers managing marine spaces. Participants’ interaction with scientists was seen as a way to legitimise data and decisions, again through personal contact and developing understanding of the processes and entities involved.

Trust in marine citizen science activities and processes, equity in accessibility to programmes, as well as capacity to learn through participation, are important factors to consider in developing long-term marine citizen science programmes (Nursey-Bray et al. 2017) and for establishing its contribution to social licence in the marine space. In developing marine citizen science to promote marine citizenship with European society, the public may be more likely to award social licence or social acceptance to marine conservation efforts by agreeing on and justifying shared community understanding on what is acceptable intervention and development in the marine realm and who is entitled to make and regulate this rules (Soma and Haggett 2015).

**Figure 1.**
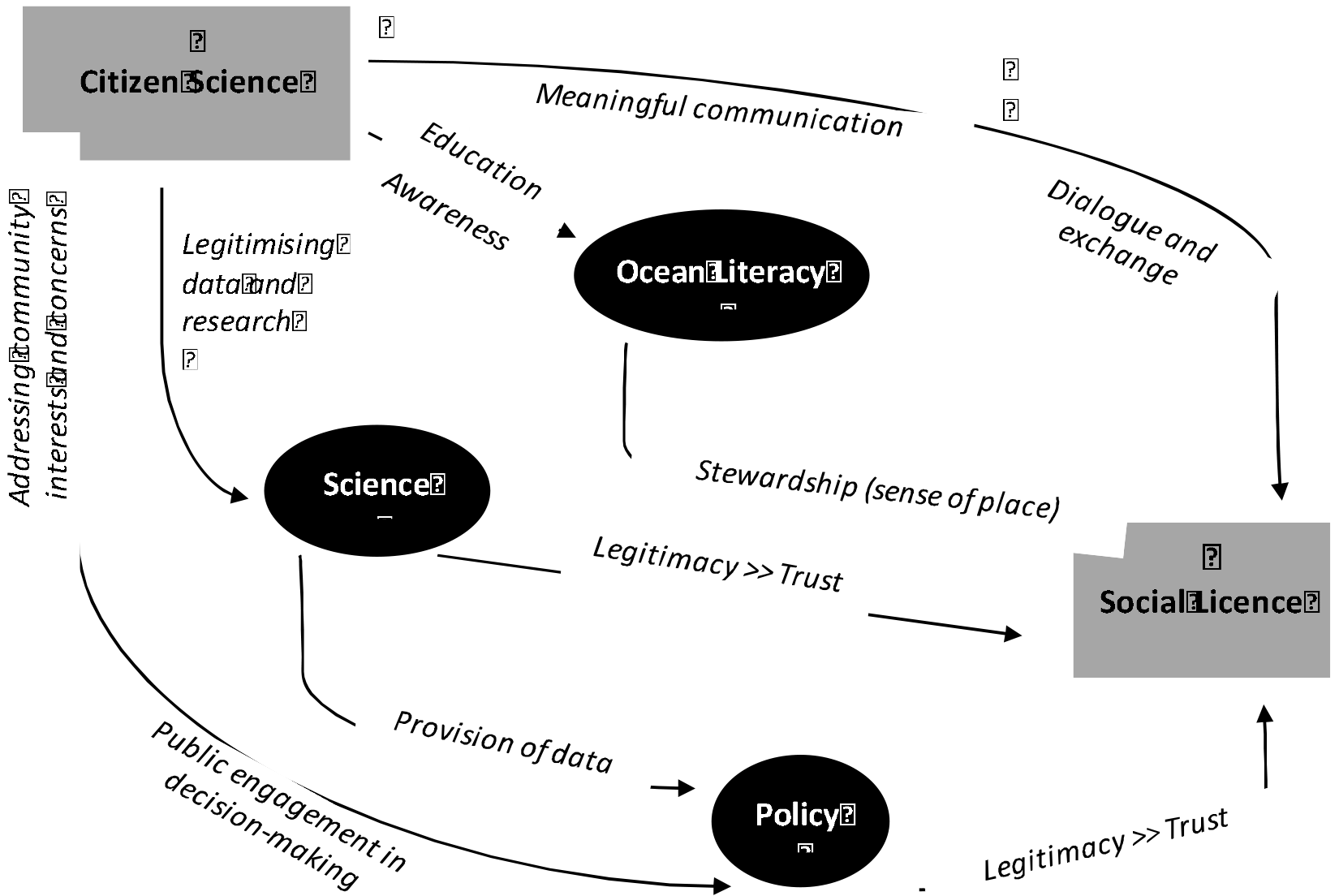
Marine citizen science provides an interface between ocean literacy and social licence

## 4. Conclusion

In this study, we adopted a largely qualitative approach to understand stakeholder perspectives and demonstrate how citizen science can play a role in enhancing social licence for marine conservation in Europe. Marine citizen science is rapidly expanding within the EU and its potential to promote ocean literacy and public engagement in policy decision making is one that requires further exploration and development (EuropeanMarineBoard 2017). Our study contributes to this research area, as we actively engaged with marine citizen science coordinators and compiled their perceptions of the utility of citizen science for this purpose, incorporating experiences and opinions from coordinators throughout Europe representing different national experiences, projects and engagement objectives. This paper reports on the European experience and we highlight that future research could conduct similar investigations on projects in regions elsewhere (including non-English speaking countries).

We have demonstrated clear linkages between citizen science and social licence (Figure 1) which are useful for exploration and application not only in a marine context, but in the terrestrial space as well. Participants in marine citizen science have the opportunity to learn and experience how science is conducted and how it contributes to conservation, decision-making and management and this can be a powerful, transformative and legitimising experience (McKinley et al. 2017). Scientific literacy can foster social empowerment and enhance public involvement in decision-making (Bela et al. 2016). Citizen science programmes provide opportunity for open discourse that is accessible to the public, (McKinley et al. 2017) and involving them in data collection and decision-making gives legitimacy to management decisions by increasing transparency (Reed 2008) and bringing stakeholders together in partnerships to contribute and develop new initiatives. Thus, marine citizen science is strategically placed to promote trust and social licence for marine conservation.

Marine citizen science can serve as a valuable platform on which to connect the public to ocean environments, but it should not be assumed that participants will automatically support ocean protection or conservation management. Generating social licence through marine citizen science requires developing meaningful relationships with participants and earning their trust through engagement, education, sharing of information, dialogue and transparency. Achieving such objectives in Europe requires planning resources and expertise, which many European marine citizen science projects do not have access to. Marine citizen science needs more and improved funding. Powerful actors such as the EU Commission (i.e. EMB) can amend this by defining and providing policy direction and support (Weiss et al. 2012).

> *‘The oceans cover most of our planet. We very much need to engage people with understanding them. At all levels, and citizen science is one of them’(B2)*.

We support growing policy calls that highlight development of marine citizen science as an imperative objective. In order to achieve these aims and to also enhance social licence for conservation, more funding opportunities will need to be made available. The costs of policy implementation associated with a lack of social licence can escalate rapidly across community, governmental, market and environmental outlays. European marine conservation requires public awareness, understanding and social licence, and marine citizen science is an ideal tool to achieve this.

## Acknowledgements

We are grateful to all the citizen science managers who offered their knowledge and expertise for this study. This research was approved by UFZ Datenschutz (Data Protection), Leipzig, Germany (23/06/2017). RK was supported by the Green Talents Awards Programme for High Potentials in Sustainable Development by the German Ministry of Education and Research (BMBF). The study also received funding from the European Union’s Horizon 2020 research and innovation programme (grant agreement no. 6417, ECOPOTENTIAL). GP was supported by an Australian Research Council Future Fellowship. The authors wish to thank the members of the Ecosystem Services research team at iDiv, Leipzig for their advice in developing this manuscript.

